# Developmental plasticity of hearing sensitivity in red-eared slider *Trachemys scripta elegans*

**DOI:** 10.1101/825968

**Authors:** Jichao Wang, Handong Li, Tongliang Wang, Bo Chen, Jianguo Cui, Haitao Shi

## Abstract

Developmental plasticity of hearing sensitivity (DPHS) has been verified in some groups of vertebrates. Turtles face a trade-off between terrestrial and aquatic hearing in different acoustic environments throughout ontogeny. However, how chelonian hearing sensitivity changes throughout ontogeny is still unclear. To verify DPHS in turtles, auditory brainstem responses (ABR) were compared using hearing thresholds and latencies in female red-eared slider (*Trachemys scripta elegans*) aged 1 week, 1 month, 1 year, and 5 years, and the results showed hearing sensitivity bandwidths of approximately 200–1100, 200–1100, 200–1300, and 200–1400 Hz, respectively. The lowest threshold sensitivity was approximately 600□Hz. Below 600 Hz, ABR threshold decreased rapidly with increasing age (1 week to 1 year), with significant differences between age groups, but no significant difference between the 1- and 5-year age groups (stimulus frequency, 200–600 Hz). Above 600 Hz, ABR threshold was the lowest in the 5-year age group. These findings show that aging was accompanied by hearing sensitivity changes, suggesting rapid, frequency-segmented development during ontogeny. This variability in hearing sensitivity differs from that reported in other vertebrates, and allows adaptation to acoustically distinct environments throughout ontogeny. Our findings further elucidate the developmental patterns of the vertebrate auditory system.

## 1. Introduction

From insects to mammals, acoustic communication is crucial for survival, successful reproduction, and many other life-history strategies [1–3]. Usually, changes in the auditory system cause changes in sensitivity during ontogeny; these can have a profound impact on an organism’s hearing [4, 5]. Moreover, the development of hearing sensitivity allows accurate and efficient acoustic communication in individuals [4].

Developmental plasticity of hearing sensitivity has been verified in fishes [6, 7], frogs [8], lizards [9], birds [10, 11], mammals [12, 13], and humans [14], suggesting that auditory processing matures with age, and that this process differs between species. Turtles, like other amphibious animals, face a trade-off between terrestrial and aquatic hearing, and their acoustic environment changes during ontogeny; they may have evolved variability in hearing capacity to adapt to complex environments [15, 16]. Although there has been considerable research into auditory system development in some groups of vertebrates, much less is known about it in chelonians. At present, we know of only three studies on the developmental plasticity of hearing sensitivity in chelonians, and these reached different conclusions [17–19]. However, these studies had small samples (n < 7), narrow age ranges, and unclear sexual categorization, factors that may affect their findings. Our previous study provided the first evidence that sexually dimorphic hearing sensitivity has evolved in turtles, with the hearing of females shows greater sensitivity [20]. It is necessary to further study ontogenetic changes in hearing sensitivity in chelonians.

Red-eared slider (*Trachemys scripta elegans*) is a semi-aquatic freshwater turtle, well adapted to living in various habitats, including rivers, streams, and even brackish water (Salinity 5.3–14.6‰) [21]. *Trachemys scripta elegans* is an important and potentially powerful model for researching hearing. Aspects of hearing, including the general ultrastructure of the auditory receptors (auditory hair cells) [22], the functional morphology of cochlear hair cell stereociliary bundles [23], the structure of the sound receiver organ (the tympanic disc) [15], habitat-related auditory plasticity [15], the response properties of auditory hair cell afferent fibers [24], the morphology of the middle-ear cavity [25], and sexually dimorphic hearing sensitivity [20], have been widely studied in this species. Those results provide an appropriate foundation and a reliable model organism to assess the developmental plasticity of hearing sensitivity in chelonians. Moreover, *T. scripta elegans* is farmed in many provinces of China; hence, sufficient numbers of experimental specimens were available.

Auditory brainstem response (ABR) measurement is a noninvasive and rapid method to measure hearing sensitivity; its use has been validated for frogs [26, 27], toads [28], and reptiles [18, 29]. Our aim was to demonstrate that aging is accompanied by changes in turtle hearing sensitivity by measuring ABR, to assess ontogenetic changes in post-hatchling-to-reproductive *T. scripta elegans* adults. To do this, we focused on the hearing sensitivity bandwidth, threshold sensitivity, and latency.

## 2. Materials and methods

### 2.1 Experimental animals

Considering that sexually dimorphic hearing sensitivity has evolved in turtles, only female *T. scripta elegans* individuals were used. We used age groups of 1 week (n = 10), 1 month (n = 11), 1 year (n = 10), and 5 years (n = 10). All animals were purchased from farms in Hainan Province, China, and maintained in standard aquaria at 20–25 °C until experiments were conducted. Body mass and carapace length are shown in Figure 1. Because we could not determine sex using external morphology, sex was determined by paraffin section of toe phalanges of 1-week and 1-month old individuals. Ages of individuals < 1 year old were determined by time since hatching. Ages of 5-year-old individuals were determined after the experiments by observing paraffin sections. Prior to electrode placement, each turtle was deeply anesthetized using a solution of 0.5% pelltobarbitalum natricum (CAS No.: 57-33-0, Xiya Reagents, Shandong, China) dissolved in 0.9% sodium chloride. The anesthetic was administered via hind limb intramuscular injection at an initial dose of 0.003 mL g−1. Additional doses (each at 20% of the initial dose) were administered in cases when the subject was not deeply anesthetized [20]. The electrophysiological experiments began after the subject showed no pain response to stimulating the hind leg muscles with forceps.

**Figure 1.**
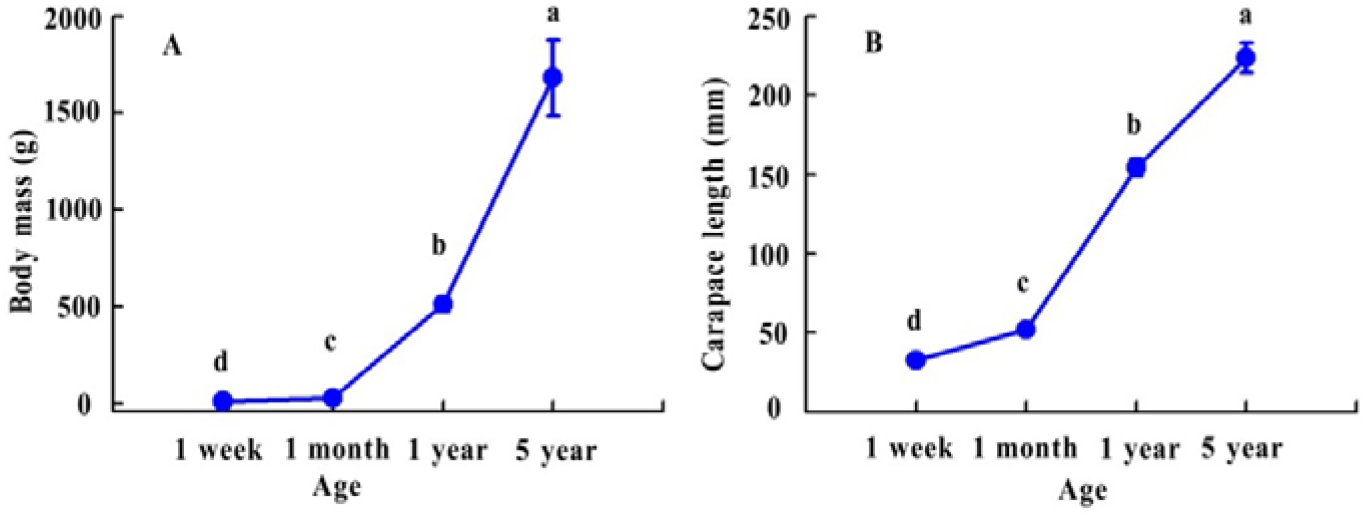
*Trachemys scripta elegans* body mass (A) and carapace length (B) varied with age. Points and error bars reflect the mean ± SD.

### 2.2 ABR measurements

#### 2.2.1 ABR procedures

ABR recordings of approximately 100 min were made using a TDT RZ6 Multi-I/O Processor, linked via fiber optic cables to a Tucker-Davis Technologies (TDT) RA4LI low-impedance digital headstage and RA4PA Medusa preamp; and analyzed using BioSig and SigGen software (Tucker-Davis Technologies, Inc., FL, USA). A portable amplified field speaker (frequency response 55–20 000 Hz, JBL GT7-6, Harman International Industries, Inc., USA) was located in a sound-proof booth lined with echo-attenuating acoustic foam, linked via fiber optic cables to a TDT RZ6 Multi-I/O Processor. Standard platinum alloy subdermal needle electrodes (27 ga, 13 mm length, Rochester Electro-Medical, Inc., Lutz, FL, USA) were inserted sub-dermally above the tympanum (recording electrode), on the top of the head under the frontal scale (reference electrode), and in the ipsilateral front leg (ground electrode), and the other end of each needle was connected to the TDT RA4LI low-impedance (<3 kΩ) digital headstage.

#### 2.2.2 Stimulus generation and presentation

Sound stimuli were generated using a TDT RZ6 Multi-I/O Processor, which directly drove the speaker, running TDT SigGen software. Each individual tone burst (9 ms duration, 2 ms rise/fall time, with a sample rate of 24 414 Hz and alternating polarity) was synthesized digitally from 0.2 kHz to 1.5 kHz in 100 Hz increments, and attenuated in 5 dB steps from 90 dB to 35 dB, and presented at a rate of 4/s. Clicks were 0.1 ms in duration with a 249 ms interstimulus interval, attenuated in 5 dB steps from 90 dB to 35 dB, and presented at a rate of 4/s. Each ABR wave represented the average response to 200 stimulus presentations. Signals from the electrodes were amplified (20×) and filtered (high pass: 30 Hz; low pass: 3 kHz; notch filtered: 50 Hz). Sound stimuli were replicated once.

#### 2.2.3 Calibration

ABR stimulus levels were calibrated in the free field using a TDT RZ6 Multi-I/O Processor and BioSigRP (Tucker-Davis Technologies, Inc., Florida, USA), linked to a sensor signal conditioner (model 480C02, PCB Piezotronics, Inc., Depew, NY, USA) with a 1/4 inch microphone (model 426B03 PCB Piezotronics, Inc., Depew, NY, USA) positioned approximately at the tip of jaw of the turtle, but when the turtle was absent. The distance between the speaker and the microphone was approximately 5 cm. The speaker repeatedly played the signal at the same rate used while recording ABRs, and simultaneously recorded the microphone signal at a sampling rate of 24 414 Hz.

#### 2.2.4 ABR thresholds and latencies define

The ABR thresholds and latencies were determined using visual inspection similar to that previously described [29]. The threshold measurement was defined as the stimulus level below which no repeatable responses could be recognized [30, 31]. In order to reduce artificial error, all turtle ABR thresholds were determined by the same experienced person. We assumed that the 80 dB level was above the ABR threshold of all turtles studied, for the stimuli used.

### 2.3 Morphological data measured

Before ABR recordings, the body mass of all specimens was recorded by an electronic balance (SI-234, Denver Instrument (Beijing) Co., Ltd.), while carapace length was measured using a Mitutoyo digital caliper (500-196-30, Mitutoyo Corp., Japan).

### 2.4 Data analysis and statistics

ABR morphologies, sensitivity thresholds, and latencies obtained from female *T. scripta elegans* in response to tone and click stimuli were sorted and analyzed using IBM SPSS 22.0 (IBM Corp., Chicago, IL, USA). Data on body mass and carapace length of different age groups were analyzed using one-way ANOVA followed by Tukey multiple comparison testing. General linear model multivariate analysis following by Tukey multiple comparison testing was used to determine the significance of differences in ABR thresholds and latencies among age groups at each stimulus frequency. Results are expressed as mean ± SD, and P < 0.05 was considered to reflect statistically significant difference.

## 3. Results

### 3.1 Morphological characteristics

Body mass was significantly influenced by age (F = 8119.45, df = 3, P < 0.001), and differed significantly among age groups (P < 0.001). Carapace length became significantly longer with age (F = 5816.28, df = 3, P < 0.001), and differed significantly among age groups (P < 0.001).

### 3.2 Hearing sensitivity bandwidth and ABR thresholds

The hearing sensitivity bandwidths were 200–1100 Hz, 200–1100 Hz, 200–1300 Hz, and 200–1400 Hz in the 1-week, 1-month, 1-year, and 5-year age groups, respectively (Fig. 2A). The greatest sensitivity frequency was about 200–900□Hz, and the lowest sensitivity measured was at about 600□Hz (Fig. 2A)

**Figure 2.**
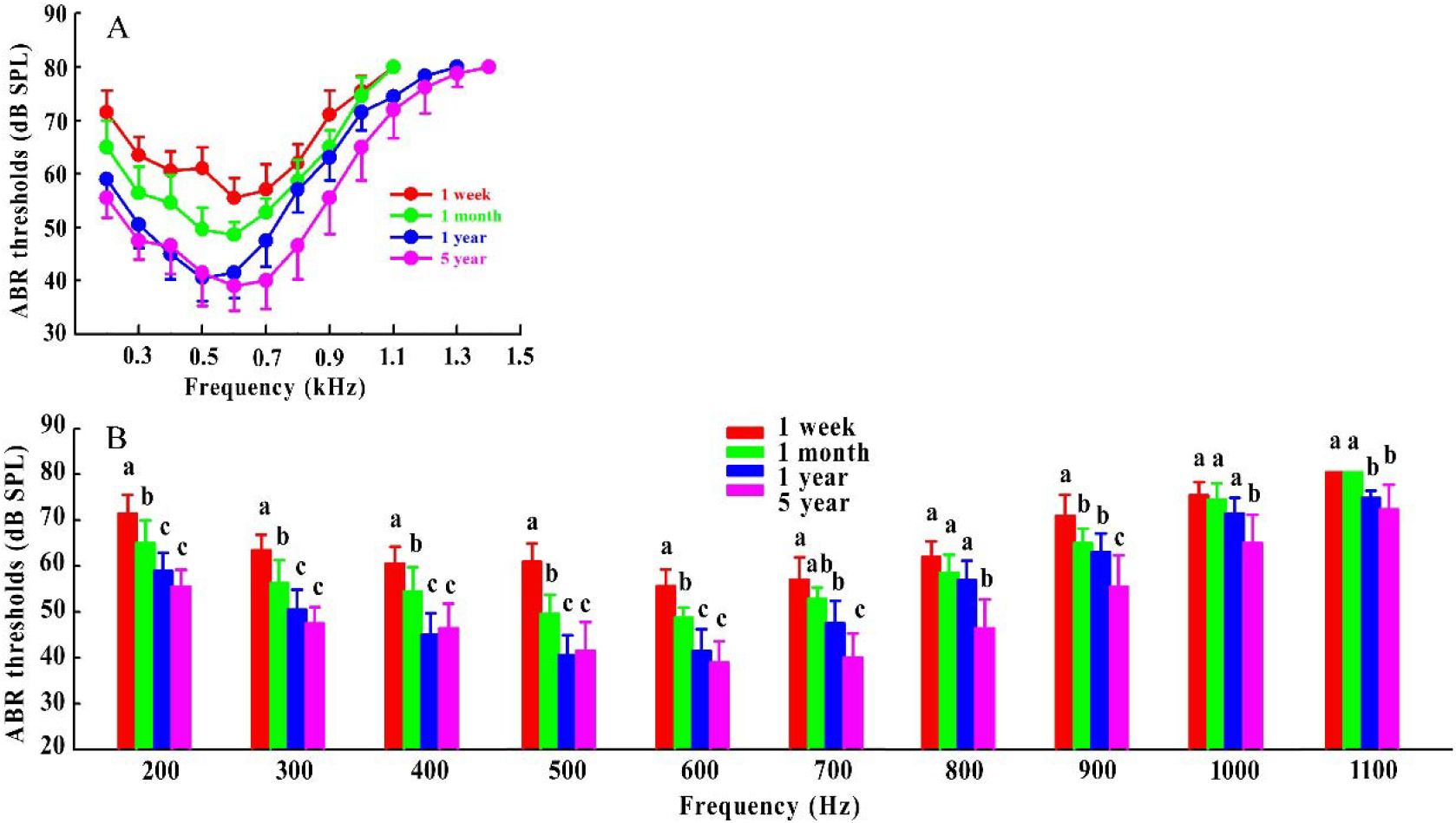
ABR threshold varied with stimulus frequency and age group in *Trachemys scripta elegans* (A); ABR threshold by stimulus frequency and age group in *T. scripta elegans* (B).

Below 600 Hz, the ABR threshold decreased rapidly with increasing age (1 week to 1 year), with significant differences among age groups, but no significant difference between the 1- and 5-year age groups, for which the stimulus frequency ranged from 200 Hz to 600 Hz (Fig. 2B). Above 600 Hz, the ABR threshold was significantly lower in the 5-year age group than in other groups (Fig. 2B).

### 3.3 ABR latency

The ABR latency for tone bursts at 75 dB was <5 ms in all age groups (Fig. 3A). The ABR latency for tone bursts was lowest in the 5-year age group (Fig. 3B).

**Figure 3.**
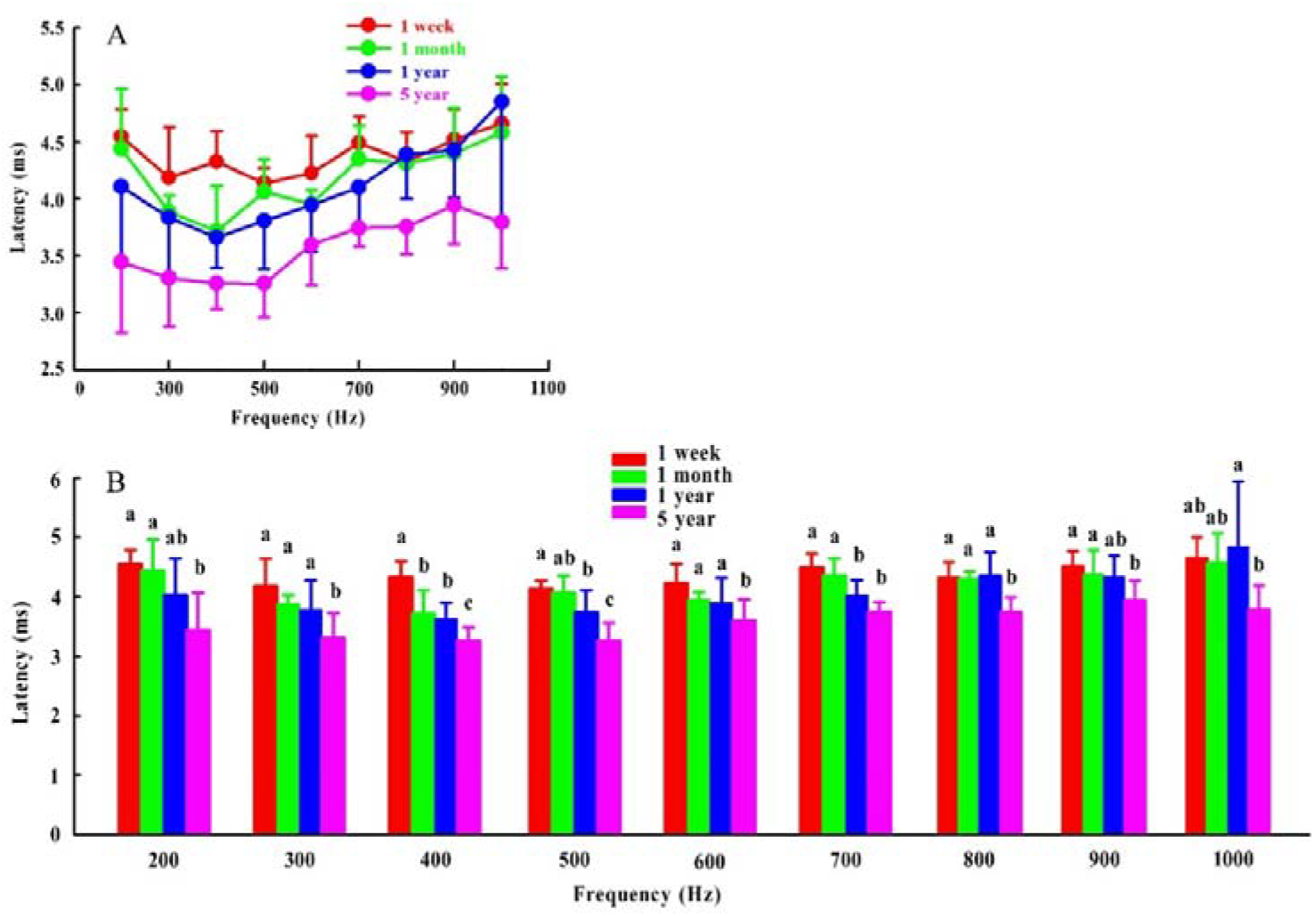
Latency against tone-burst frequency at 75 dB, by age group, in *Trachemys scripta elegans* (A); Latency by stimulus frequency and age group in *T. scripta elegans* (B).

## 4. Discussion

Herpetologists have long considered turtles and tortoises to be “the silent group”, neither vocalizing nor hearing particularly well [32, 33]. However, the hearing sensitivity of chelonians has recently begun to attract attention, and behavioral and electrophysiological studies have revealed that turtles and tortoises are low-frequency specialists (usually <1000 Hz) [16, 34, 35]. We found that the turtles could not hear frequencies >1400 Hz, and that the upper limit of the hearing sensitivity bandwidth shifted higher during ontogeny.

Although habitat-related auditory plasticity has been reported for *T. scripta elegans* [15], little is known about the development of the auditory system in this group. To our knowledge, ours is the first study to demonstrate developmental plasticity of peripheral hearing sensitivity from the post-hatchling to reproductive adult stage in chelonians. We found that the ABR threshold was significantly lower in the 1-week to 1-year age groups than in the 5-year age group. Remarkably, frequency sensitivity was developmentally segmented by 600 Hz: below this, the ABR threshold did not differ among the 1-to 5-year age groups; above this, it was significantly lower in the 1-year than 5-year age group. We thus conclude that ontogenetic hearing sensitivity development is rapid and frequency-segmented in *T. scripta elegans*. Our findings differ somewhat from those reported elsewhere. For instance, 3-year-old Hawksbill turtles (*Eretmochelys imbricata*) were sensitive to a wider frequency range and exhibited a larger amplitude response than the 2-year-olds [18]; loggerhead sea turtles (*Caretta caretta*) exhibited little difference in threshold sensitivity and frequency bandwidth throughout ontogeny [19]; and in subadult Green turtles (*Chelonia mydas*), smaller individuals had a wider hearing sensitivity bandwidth (100–800□Hz) than larger individuals (100–500□Hz) [17]. Our study improves on earlier studies, potentially providing more reliable results, because we used a larger age range, and, more importantly, we first determined the sex of individuals.

In the Bicolor damselfish (*Pomacentrus partitus*) the auditory thresholds decrease exponentially with increasing age, rapidly approaching adult levels [36]. Hearing sensitivity changes only slightly during growth of the Lusitanian toadfish (*Halobatrachus didactylus*) [37]. In budgerigars (*Melopsittacus undulatus*), hearing is poor at hatching, and thresholds improve markedly in the first week; by 1 week before fledging, ABR audiograms of young budgerigars are very similar to those of adult birds [11]. In some mammals, hearing sensitivity is weak at birth and gradually develops during the first weeks of life [38, 39], although other species, such as *Phyllostomus discolor* bats [13] and humans [40], have a well-developed auditory system at birth and even before birth. Our findings for *T. scripta elegans*, which include hysteresis in ontogenetic auditory development, therefore differ from those reported in other vertebrates. Our results further suggest that the time required to achieve a final level of auditory maturation varies among species.

Three possible mechanisms of developmental plasticity of peripheral hearing sensitivity in vertebrates have been reported. First, in frogs, the size of the tympanic membrane may be linked to differences in hearing sensitivity [41], and in some species of lizards, increased body size (or age) is accompanied by functional changes in the auditory periphery [9]. For *T. scripta elegans*, the tympanic disc is the key sound receiver [15]. However, we have found that the size of the tympanic membrane is not related to sexual dimorphism in hearing sensitivity in turtles [20]; thus, growth of the tympanic membrane during ontogeny may not explain the developmental plasticity of hearing sensitivity that we report herein. Second, age-related changes in middle-ear sound conduction occur [9]; once the structural development of the middle ear is complete, adult-like sound conduction is exhibited [9, 42–44]. Research into habitat-related plasticity of hearing sensitivity has shown that *T. scripta elegans* is more sensitive to sound underwater than in the air, and that this is related to the large size of the middle ear [15]. Consequently, age-related changes in middle-ear sound conduction may also contribute to age-related alterations in threshold sensitivity in *T. scripta elegans*. Third, the sensory epithelium of the cochlear receptor organ may increase in size throughout life. In frogs and fishes, the area of auditory receptor epithelium increases with age, and cochlear growth is accompanied by an increase in the number of hair cells on the sensory surface [9, 45–47]. This suggests that the number of hair cells is related to developmental plasticity of hearing sensitivity in *T. scripta elegans*. Future morphological and anatomical research should address these questions.

Adult female turtles spend more time on land during the reproductive period, when laying and incubating; their improved hearing may enable them to adapt to the complex terrestrial environment. Hence, the rapid and frequency-segmented development of the chelonian auditory system during ontogeny reflects adaptation to acoustically distinct environments, and prepares the adults for reproduction.

In conclusion, we found that there was rapid and frequency-segmented development of the auditory system during ontogeny in *T. scripta elegans*. This differs from what has been reported for other vertebrates. Our findings provide greater clarity on the patterns of development of the vertebrate auditory system. It is still unknown whether age-related changes in middle-ear structures and in the auditory receptor epithelium lead to developmental plasticity of peripheral hearing sensitivity in this species. Further, as sexually dimorphic hearing sensitivity has been found in turtles, it is worth investigating whether they also exhibit sexual dimorphism in developmental plasticity of hearing sensitivity. Future morphological and anatomical studies should address these questions.

## Acknowledgements

We would like to thank Yao Sun, Chunhua Zhou, and Xintong Li for their assistance during the study.

